# Eiger/TNFα-mediated Dilp8 and ROS production coordinate intra-organ growth in *Drosophila*

**DOI:** 10.1101/606707

**Authors:** Juan A. Sanchez, Duarte Mesquita, María C. Ingaramo, Federico Ariel, Marco Milán, Andrés Dekanty

**Affiliations:** Instituto de Agrobiotecnología del Litoral, Consejo Nacional de Investigaciones Científicas y Técnicas (CONICET); Facultad de Bioquímica y Ciencias Biológicas, Universidad Nacional del Litoral (UNL), Santa Fe, 3000, Argentina; Institute for Research in Biomedicine (IRB Barcelona), The Barcelona Institute of Science and Technology; Institucio Catalana de Recerca i Estudis Avançats (ICREA), Barcelona, Spain

**Author notes:** Correspondence to: Andres Dekanty, int. 5022; Marco Milán, /34 93 4037109.

**Keywords:** *Drosophila*, organ growth, coordination, p53, JNK, Eiger/TNFα, Dilp8, ROS

## Abstract

Coordinated intra- and inter-organ growth is essential during animal development to generate individuals of proper size and shape. The *Drosophila* wing has been a valuable model system to reveal the existence of a stress response mechanism mediated by Drosophila p53 (Dmp53) and involved in the coordination of tissue growth between adjacent cell populations upon targeted reduction of growth rates. Here we present evidence that a two-step molecular mechanism is being used by Dmp53 to reduce in a non-autonomous manner growth and proliferation in adjacent cell populations. First, Dmp53-mediated transcriptional induction of *Drosophila* TNFα ligand Eiger leads to cell autonomous activation of JNK. Second, two different signaling events downstream of the Eiger/JNK axis are induced in the growth depleted territory in order to independently regulate tissue size and cell number in adjacent cell populations. Whereas expression of the systemic hormone dILP8 coordinates intra-organ growth and final tissue size, induction of Reactive Oxygen Species downstream of Eiger/JNK and, as a consequence of apoptosis induction, acts non-cell-autonomously to regulate proliferation rates of adjacent epithelial cells. Our results unravel how local and systemic signals can act concertedly to coordinate growth and proliferation within an organ in order to generate well-proportioned organs and functionally integrated adults.

**Author Summary:** Coordination of growth between the different parts of a given developing organ is an absolute requirement for the generation of functionally integrated structures during animal development. Although this question has fascinated biologists for centuries, the responsible molecular mechanisms have remained so far unknown. In this work, we have used the developing wing primordium of Drosophila to identify the molecular mechanisms and signaling molecules mediating communication between adjacent cell populations upon targeted reduction in growth rates. We first present evidence that activation of Drosophila p53 in the growth-depleted territory induces expression of the fly TNF ligand Eiger which cell autonomously activates the JNK stress signaling pathway. While JNK-dependent expression of the systemic hormone dILP8 reduces growth and final size of the adjacent territories, production of Reactive Oxygen Species downstream of JNK and the apoptotic machinery act locally to regulate proliferation rates in adjacent epithelial cells. Our data reveal how signals acting locally or systemically can regulate cell proliferation and growth independently to accomplish coordination in tissue size and cell number among different parts of an organ in order to give rise to well-proportioned adult structures.

**HIGHLIGHTS:**

✓ Dmp53-dependent Eiger expression is required to coordinate intra-organ growth
✓ Eiger acts through its receptor Grindelwald and JNK signaling to reduce growth and proliferation rates in a non-cell-autonomous manner
✓ Eiger/JNK-dependent Dilp8 expression coordinates intra-organ growth but not proliferation
✓ Eiger/JNK activation triggers ROS production
✓ ROS act non-cell-autonomously to regulate proliferation of adjacent epithelial cells.

## INTRODUCTION

Coordinated tissue growth is essential for the generation of functionally integrated organs during animal development as well as for tissue homeostasis during adult life. Even though a broad range of genes and pathways regulating growth has been uncovered, the mechanisms by which cells within the same tissue are able to respond to stress in a coordinated manner and maintain tissue homeostasis remain less well understood.

The p53 tumor suppressor regulates the response of mammalian cells to stress through direct transcriptional activation of specific target genes involved in cell cycle arrest, DNA repair and apoptosis [1]. Recently, several non-cell-autonomous functions of p53 have become relevant in tissue homeostasis as well as tumor suppression [2,3]. For instance, it has been reported that activation of p53 in stromal fibroblast promotes an antitumor microenvironment by influencing survival and spreading of adjacent tumor cells [4–6]. Consistently, reconstituted MCF7 tumors containing p53-deficient fibroblasts developed faster and were more aggressive than their counterparts carrying wild-type fibroblasts [7]. In addition, modulation in the secretion of inflammatory cytokines by tumor-mediated activation of p53 has been shown to limit tissue damage [8], inhibit adjacent epithelial cell transformation [4] and promote macrophage-mediated clearance of tumor and apoptotic cells [9,10].

Studies on p53 coordinating cell growth and proliferation have benefited from genetically tractable model systems. The single *Drosophila* orthologue of mammalian p53 (Dmp53) has been demonstrated to be essential for tissue and metabolic homeostasis [2]. In addition to conserved functions promoting apoptosis and cell-cycle arrest upon stress, Dmp53 participates in non-cell-autonomous responses regulating apoptosis-induced proliferation [11–13], cell competition [14,15] and adaptive responses to nutrient stress at the organismal level [16]. Previous studies using *Drosophila* wing as a model system demonstrated that adjacent cell populations within the organ are able to grow in a coordinated manner buffering local growth rates variations to maintain tissue homeostasis and produce well-proportioned wings [17,18]. Depletion of growth promoting genes or disruption of the protein biosynthetic machinery in specific compartments of the developing wing primordium non-cell-autonomously reduces the size of adjacent, unperturbed territories [17,19]. Activation of Dmp53 in the slow growing compartment is required for proper intra-organ growth coordination as depletion of Dmp53 uncouples growth of adjacent cell populations and ultimately gives rise to wings with wrong proportions [17]. Several molecular mechanisms have been proposed to underlie Dmp53-dependent coordination of growth, either involving a tissue local response [17] or systemic mechanisms coordinating both intra- and inter-organ growth [19–21].

In this study, we have identified Eiger, the unique member of TNF superfamily of ligands in *Drosophila*[22,23], as a direct Dmp53 downstream effector in coordinating intra-organ growth. We demonstrate that Eiger acts through Grindelwald (Grnd) and JNK signaling to reduce growth and proliferation rates of adjacent cell populations. Inactivation of Eiger or JNK signaling in slow growing compartments disrupts coordination of growth among adjacent tissue domains resulting in unproportioned adult wings. We further demonstrate that regulation of tissue growth and proliferation rates by Dmp53 can be uncoupled and independently regulated by two different mechanisms downstream of Eiger-JNK signaling. Whereas dILP8 expression is required to coordinate intra-organ growth and final tissue size, ROS production downstream of Eiger/JNK and as a consequence of apoptosis induction acts non-cell-autonomously to regulate proliferation rates of adjacent epithelial cells. Taken together, our results show how tissue local mechanisms along with systemic responses can act together to coordinate growth and proliferation between the different parts of an organ.

## RESULTS

### Dmp53 target genes are upregulated upon growth stress

The *Drosophila* wing imaginal disc is a highly proliferative epithelium that grows extensively during larval development and provides an ideal model to study intra-organ growth coordination [24,25]. We made use of the Gal4/UAS system to direct expression of a cold sensitive version of type 2 ribosome-inactivating protein Ricin-A (RA^CS^) or an RNAi for the growth promoting transcription factor dMyc (*dmyc*^RNAi^) to specific domains in the wing primordium. This allows a direct comparison between cells expressing RA^CS^ or *dmyc*^RNAi^ and adjacent wild-type cells of the opposite compartment. Consistent with previous reports, targeted expression of RA^CS^ or *dmyc*^RNAi^ to the posterior (P) or anterior (A) compartments of the wing disc reduces size of both transgene-expressing and adjacent wild-type territories (Figure 1A-C; [17]). Along with size decrease, a non-autonomous reduction in proliferation rates and final cell number was also observed as assessed by measurements of BrdU incorporation (Figure 1D-F) and cell density in adult wings (Figure 1G-H; [17]).

**Figure 1:**
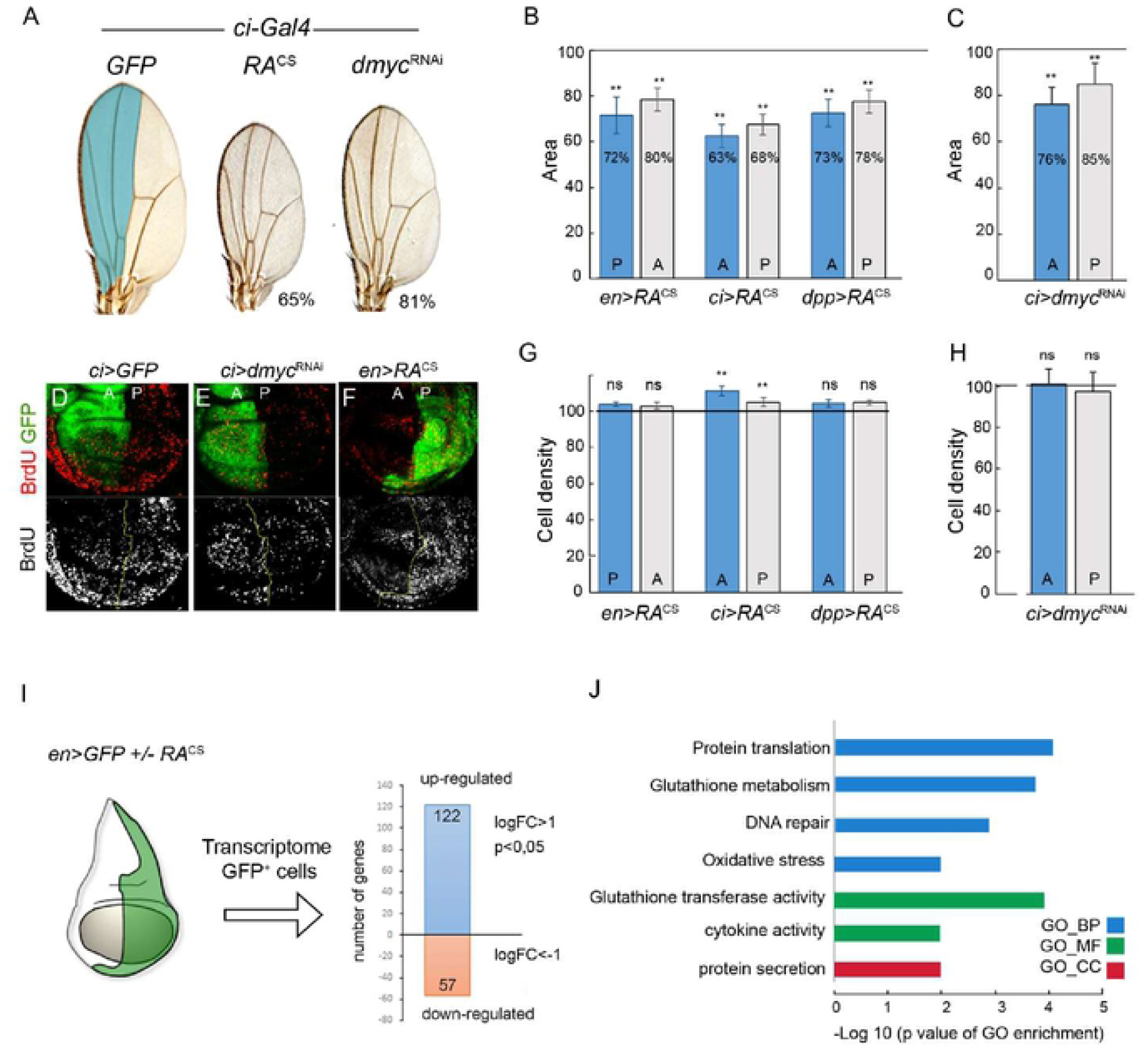
Coordinated intra-organ growth involves upregulation of Dmp53 target genes. (A) Representative adult wings from individuals expressing GFP, *dmyc*^RNAi^ or RA^CS^ under the control of *ci-Gal4*, which drives expression to the anterior compartment of wing imaginal discs (showed in light blue). Numbers indicate total wing area represented as a percentage of GFP expressing wings. (B-C, G-H) Histograms plotting normalized area (B-C) or cell density values (G-H) of the anterior (A) and posterior (P) compartments of adult wings from individuals expressing RA^CS^ or *dmyc*^RNAi^ with the indicated Gal4 drivers. Blue bars indicate the transgene-expressing compartment and grey bars indicate the adjacent wild-type compartment. Expression of RA^CS^ or *dmyc*^RNAi^ significantly reduced the size of both transgene-expressing and non-expressing domains. Horizontal line shows the size or cell density values of the normalized control GFP expressing wings. ** p<0.01. (D-F) BrdU incorporation assay in larval wing discs from individuals expressing GFP along with the indicated transgenes under the control of *ci-Gal4* or *en-Gal4*. (I) Transcriptome analysis of RNA samples obtained from posterior compartment (GFP positive cells) of wing imaginal discs from individuals expressing GFP or GFP plus RA^CS^ with *en-Gal4*. 179 genes were differentially expressed between RA^CS^ expressing and wild type cells (−1≥fc≥1; p≤0.05). (J) Gene ontology (GO) analysis showed highly enriched GO terms amongst genes upregulated in RA-expressing wing discs. Log 10 of p-value are shown. GO-BP, biological processes. GO-MF, molecular function. GO-CC, cellular component.

In order to identify novel molecules and pathways involved in coordinating intra-organ growth, we performed differential gene expression analysis of wing imaginal discs expressing RA^CS^ (*en-Gal4; UAS-RA, UAS-GFP*) and control wing discs expressing GFP (*en-Gal4;UAS-GFP*). Transcriptome analysis of RNA samples from P compartment cells (GFP positive cells) of larvae of the indicated genotypes identified 179 genes that were differentially expressed (Figure 1I and Table S1). Quantitative real-time RT-PCR (qRT-PCR) measurements showing consistent upregulation of several of these genes in cells expressing either RA^CS^ or *dmyc*^RNAi^ strongly suggest both perturbations elicit a common cellular response (Figure S1). Gene ontology (GO) analysis comprising the 179 identified genes revealed an enrichment in biological processes associated with cellular responses to DNA damage, oxidation-reduction processes, glutathione metabolism, cytokine signaling and extracellular proteins (Figure 1J). In addition, 57 genes previously described as p53 targets [26,27] were specifically up-regulated upon RA^CS^ expression (Figure S1). Together, these results validate previous reports and strongly support the proposed role for Dmp53 in coordination of tissue growth.

### Eiger/TNFα is required to coordinate intra-organ growth downstream of Dmp53

Next, we looked for upregulated secreted signaling molecules in RA^CS^-expressing cells that could mediate tissue non-autonomous responses. GO analysis recognized a group of 19 genes coding for extracellular proteins (Figure 1H, Figure S2 and Table S3). To test whether these molecules and others were actually relevant for intra-organ growth coordination, we decided to perform an *in vivo* loss-of-function screen using *Drosophila* transgenic RNAi lines (Table S2). Genes encoding for potential secreted factors were specifically silenced in the RA^CS^-expressing domain and screened for their capacity to affect the size of the adjacent compartment in adult wings. Notably, expression of *eiger/TNFα*^RNAi^ partially rescued the non-autonomous reduction in tissue size caused by RA expression resulting in adult wings with not well-proportioned compartments (Figure 2A-B). We further confirmed the role of Eiger in non-autonomous regulation of tissue size by using an independent RNAi line (*UAS-egr*^IR^) (Igaki et al., 2002) and *eiger* homozygous mutants (Figure 2B and Figure S2).

**Figure 2:**
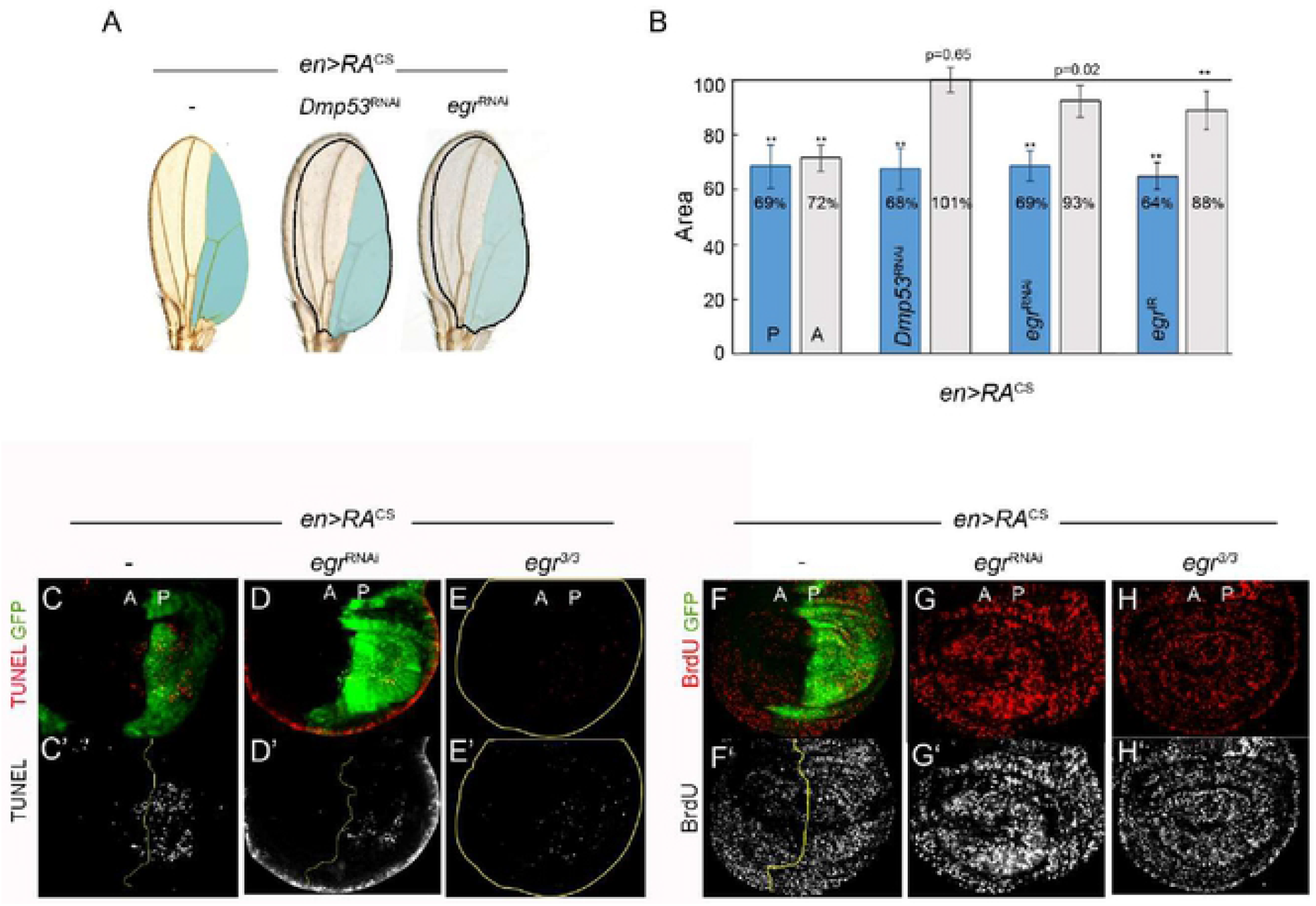
Eiger/TNFα non-autonomously regulates growth and proliferation rates. (A) Representative adult wings from individuals expressing the indicated transgenes in posterior compartment (blue) under the control of *en-Gal4*. The expected size of the neighboring domain to give rise to a well-proportioned adult wing is depicted. (B) Histogram plotting normalized area of the posterior (P; blue bars) and anterior (A; grey bars) compartments of adult wings from individuals expressing the indicated transgenes with *en-Gal4*. Expression of *Dmp53*^RNAi^, *eiger*^RNAi^ or *eiger*^IR^ reverted the non-autonomous reduction in tissue size caused by RA expression. ** p<0.01. (C-H) Wing discs expressing GFP along with RA^CS^ in the P compartment by using *en-Gal4* in different genetic backgrounds (*wildtype* (C,F); co-expressing *eiger*^RNAi^ (D,G) or *eiger^3/3^* (E,H)) and labeled to visualize GFP, TUNEL and BrdU incorporation.

It has been previously reported that Eiger participates in apoptosis-induced apoptosis (AiA), a process by which dying cells in one compartment induce apoptosis in the adjacent one [28]. Interestingly, RA^CS^ expression in *Drosophila* wing discs elicited apoptosis in both transgene-expressing and adjacent wild-type territories (Figure 2C; [17,29]). We then assessed Eiger functional requirement for apoptosis induction and found that RA^CS^-induced apoptosis was strongly suppressed in both wing disc compartments when depleting Eiger (Figure 2D-E). Whereas apoptosis is not required for wing size reduction, it plays an essential role in non-autonomous decrease of proliferation rates [17]. We then investigated whether Eiger is also responsible for reducing proliferation rates upon RA^CS^ expression. Interestingly, expression of *egr*^RNAi^ largely rescued the non-autonomous effects of RA on BrdU incorporation levels and number of mitotic PH3-positive cells (Figure 2F-G and Figure S2). Similar results were observed in homozygous *egr^3^* mutant animals (Figure 2H). Together, these evidences support a fundamental role of Eiger in regulating the size and cell number of adjacent tissue domains.

Supporting transcriptome data, qRT-PCR assays showed increased *egr* transcript levels in RA^CS^ and *dmyc*^RNAi^ expressing cells (Figure 3D and S1). We then analyzed *egr* expression by RNA *in situ* hybridization and found a strong increase in *egr* transcript levels in the RA^CS^- or *dmyc*^RNAi^-expressing wing disc territory (Figure 3A-C). We confirmed these findings using *in vivo* reporters of both *egr* transcription (*egr*-lacZ; [30]) and Eiger protein expression and localization (Eiger-GFP; [30,31]). *dmyc*^RNAi^-expressing cells showed strong activation of egr-lacZ (Figure 3E-F) and displayed increased Eiger-GFP levels (Figure S3), therefore indicating that Eiger is specifically and cell-autonomously induced in wing discs upon RA^CS^- or *dmyc*^RNAi^-expression.

**Figure 3:**
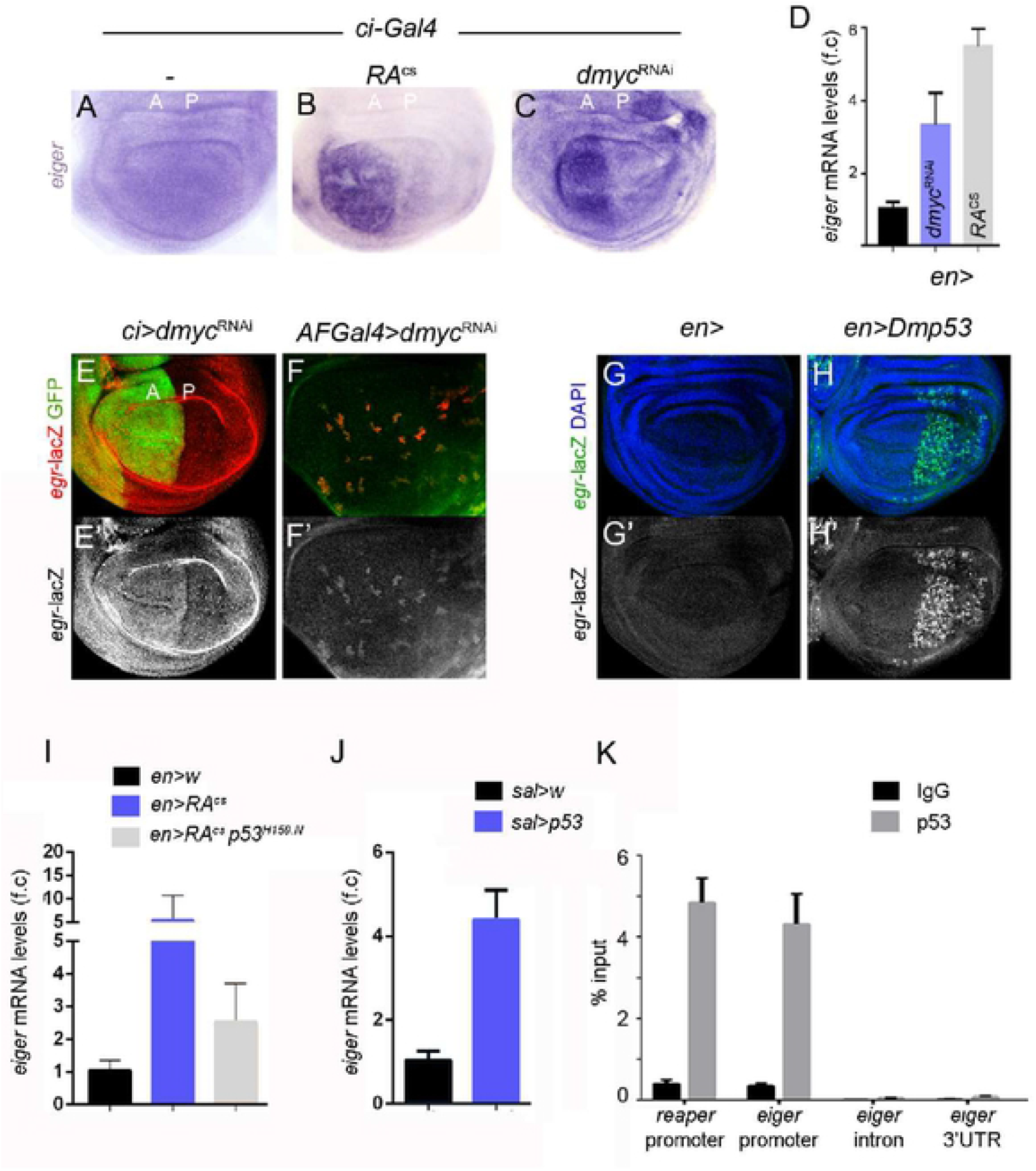
Dmp53-dependent Eiger expression in slow growing cell populations. (A-C) *In situ* hybridization to visualize *eiger* transcript levels in wing discs from individuals expressing GFP (A), RA^CS^ (B) or *dmyc*^RNAi^ (C) under the control of *ci-Gal4*. (D) qRT-PCR showing *eiger* mRNA levels in wing discs expressing *RA^CS^* or *dmyc^RNAi^* with *en-Gal4*. Results are expressed as fold induction relative to control wing discs. (E-F) Wing discs carrying *egr*-lacZ transcriptional reporter and stained to visualize GFP (green) and β-gal (red in E-F; grey in E’-F’) protein expression. *dmyc*^RNAi^ is expressed in anterior cells with *ci-Gal4* (E) or in clones using *Actin-Flipout-Gal4* (F). Note increased levels of egr-lacZ in dMyc-depleted cells. (GH) Wing imaginal discs from controls (G) or expressing Dmp53 (H) under the control of *en-Gal4; tub-Gal80^ts^* and stained to visualize egr-lacZ (green or grey) and DAPI (blue). (I-J) qRT-PCR showing *eiger* mRNA levels in wing discs expressing the indicated transgenes under the control of *en-Gal4* (I) or *sal-Gal4;tub-Gal80^ts^* (J). Results are expressed as fold induction with respect to control wing discs. (K) ChIP assays from larvae expressing Dmp53 (*sal-Gal4/+;tub-Gal80^ts^/UAS.Dmp53*) using anti-p53 antibodies or unrelated IgG (control) followed by qPCR for a region overlapping predicted p53-binding elements at *eiger* and *reaper* promoters. *eiger* intronic or 3’UTR regions were used as negative controls. Data (qPCR) are mean ± s.d.

We next asked whether Dmp53 was responsible for the expression of *egr* in RA^CS^-expressing wing discs. Interestingly, expression of a dominant negative version of Dmp53 lacking DNA-binding activity (Dmp53^N.159H^) partially reduced RA-induced *egr* expression as well as induction of *corp*, a known Dmp53 target gene (Figure 3I and S3). We next tested whether Dmp53 overexpression was sufficient to activate *egr* transcription. Conditional expression of Dmp53 in wing imaginal disc using Gal4/Gal80^ts^ system showed a 4-fold increase in *egr* mRNA levels (Figure 3J) and strongly induced egr-lacZ expression (Figure 3G-H). Consistent with recent reports showing Dmp53 binding to *egr* locus [32], chromatin immunoprecipitation (ChIP) assays showed that Dmp53 binds to *egr* and *reaper* predicted binding sites with similar efficiency (Figure 3K). Altogether, these results are consistent with Dmp53-activating *egr* gene expression through a conserved p53 binding site in *egr* promoter region.

### Eiger-induced JNK signaling is required for non-cell-autonomous reduction of growth and proliferation rates

The Jun N-terminal Kinase (JNK) cascade is a conserved stress response pathway and an important regulator of tissue growth, proliferation and apoptosis [23,33–37]. Our transcriptome analysis indicated that RA expression activates JNK signaling as we observed the induction of JNK activators such as *eiger*, *gadd45* and *traf4*, and JNK target genes such as the JNK-phosphatase *puckered* (*puc*), *Matrix Metalloproteinase 1* (*MMP1*), *insulin-like-peptide 8* (*dilp8*) and *PDGF- and VEGF-related factor 1* (*Pvf1*)(Figure S1 and Table S3). Consistently, we observed activation of the JNK reporter puc-lacZ in wing discs expressing RA^CS^ (Figure 4A) and a significant increase in MMP1 protein levels when expressing *dmyc*^RNAi^ (Figure 4B-D). MMP1 expression was completely blocked in *hep^r75^* heterozygous animals (Hemipterous; JNK kinase homologue in *Drosophila*) or upon expression of a dominant negative version of Basket (Bsk^DN^; JNK homologue in *Drosophila*) (Figure 4F-G). To test whether JNK pathway is activated downstream of Egr in slow growing tissues, we depleted Egr or TNFα receptor homologue Grindewald (Grnd) [38] and measured MMP1 protein levels. As shown in Figure 4H-I, MMP1 ectopic expression caused by *dmyc* knockdown was entirely reverted by expression of either *egr*^RNAi^ or grnd^RNAi^. Additionally, expression of Dmp53^N.159H^ largely impaired JNK activation caused by *dmyc*^RNAi^ or RA^CS^ expression (Figure 4E and Figure S4). Collectively, these results indicate that Dmp53 activation in growth impaired territories drives Eiger-dependent activation of JNK signaling pathway.

**Figure 4:**
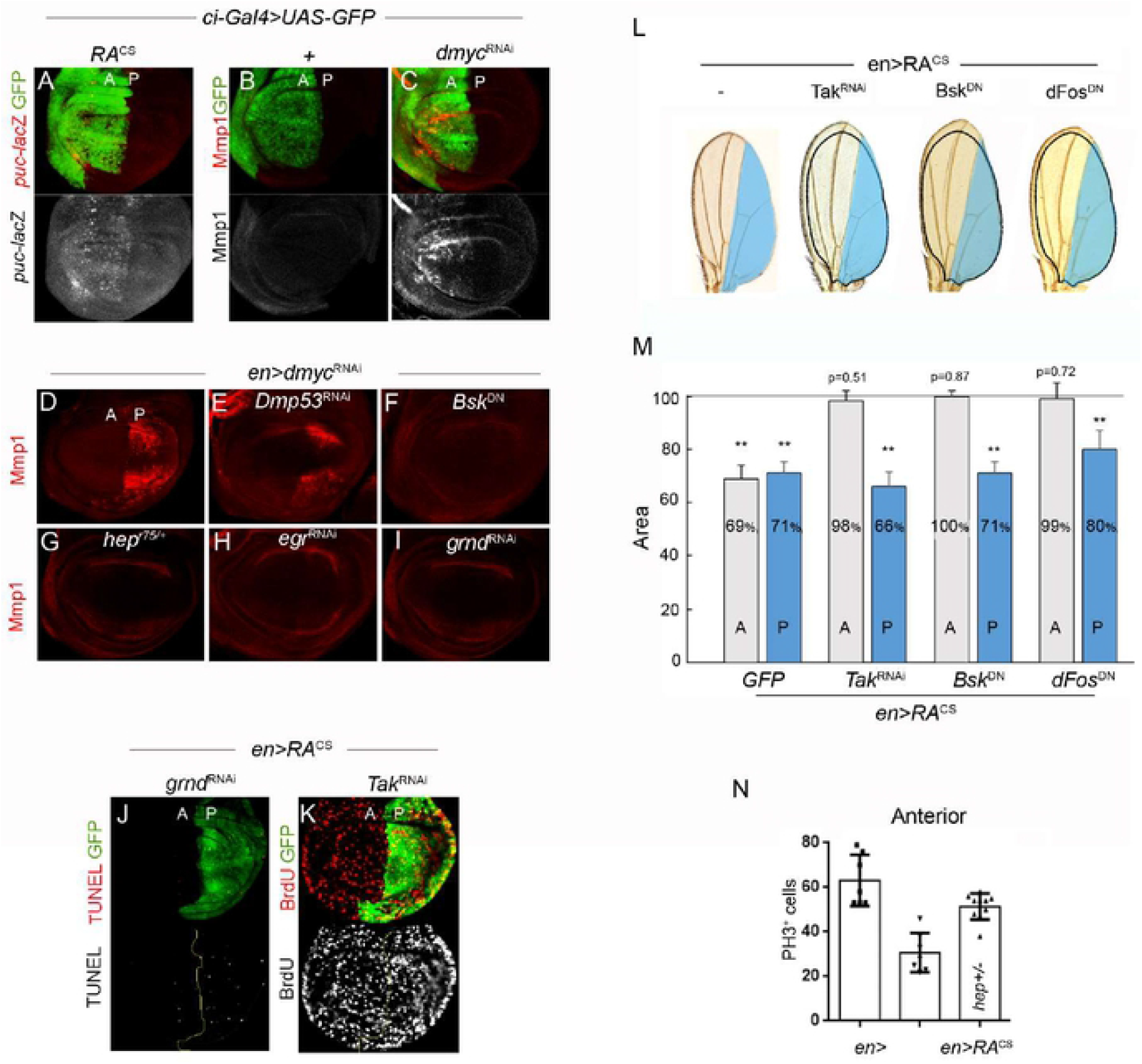
JNK signaling non-autonomously regulates growth and proliferation rates. (A-C) Wing imaginal discs from individuals expressing GFP, RA^CS^ or *dmyc*^RNAi^ and stained to visualize GFP (A-C), *puckered-lacZ* (A) and MMP1 (B-C). (D-I) Wing discs labeled to visualize MMP1 protein expression from individuals expressing *dmyc*^RNAi^ alone or in combination with *Dmp53*^RNAi^ (E), Bsk^DN^ (F), *hep^r75/+^* (G), egr^RNAi^ (H) and grnd^RNAi^ (I). (J-K) Wing imaginal discs from individuals expressing RA^CS^ along with the indicated transgenes under the control of *en-Gal4* and stained to visualize TUNEL (J) and BrdU incorporation (K). (L) Representative adult wings from individuals expressing the indicated transgenes in posterior compartment (blue) under the control of *en-Gal4*. The expected size of the neighboring domain to give rise to a well-proportioned adult wing is depicted. (M) Histogram plotting normalized area of the posterior (P; blue bars) and anterior (A; grey bars) compartments of adult wings from individuals expressing RA along with the indicated transgenes with *en-Gal4*. Blocking JNK pathway by co-expression of *Tak*^RNAi^, *Bsk*^DN^ or *Fos*^DN^ totally reverted the non-autonomous reduction in tissue size caused by RA expression. ** p<0.01. (N) Histogram plotting PH3 positive cells in the anterior (A) compartment of wing imaginal discs from individuals expressing the indicated transgenes under the control of *en-Gal4*.

The fact that MMP1 and *puckered* expression were restricted to the transgene expressing cell population suggests that Eiger/JNK signaling is mediating an autonomous apoptotic response. Indeed, depletion of *Grnd* or expression of Bsk^DN^ markedly reduced the number of TUNEL positive cells caused by RA^CS^ expression (Figure 4J and data not shown). Next, we asked whether, in addition to the expected role in apoptosis, JNK signaling might contribute to the non-autonomous response of the tissue. To address this, we inhibited JNK signaling specifically in RA^cs^-expressing compartment and analyzed proliferation rates in adjacent cell populations. Interestingly, when activation of JNK pathway was reduced, either by using *Tak1*^RNAi^ or *hep^r75^* mutants, the non-autonomous effects of RA^CS^ expression on the levels of BrdU incorporation and on the number of mitotic PH3-positive cells were largely rescued (Figure 4K, N). We next analyzed the resulting adult wings when the activation of JNK pathway was impaired. Notably, the non-autonomous reduction in tissue size caused by RA expression was completely rescued when co-expressing *Tak1*^RNAi^ or *Bsk*^DN^ (Figure 4L-M). Identical results were obtained with a dominant negative form of dFos, a transcription factor acting downstream of JNK signaling (Figure 4L-M). Together, these results indicate that, upon growth depletion, JNK exerts an essential non-autonomous role in reducing the size of adjacent cell populations.

### Dilp8 is required to coordinate intra-organ growth downstream of Eiger/JNK signaling

Recently, *Drosophila* insulin-like peptide 8 (Dilp8) has been identified as a signaling molecule produced by imaginal discs in response to tissue damage [39,40]. Dilp8 production by slow growing or damaged tissues results in activation of neuronal receptor Lgr3 which in turn downregulates ecdysone synthesis therefore delaying metamorphosis and systemically reducing growth [39–43]. Interestingly, we observed a strong induction of *dilp8* transcript levels upon RA^CS^ or *dmyc*^RNAi^ expression (Figure 5A, Figure S1 and Table S1). Similarly, we observed induction of a dilp8^MI00727^ protein trap reporter ([40]; hereafter Dilp8-GFP) in wing discs expressing RA^CS^ (Figure 5C-D). JNK signaling has been shown to induce Dilp8 expression in response to various types of stress [39,44,45]. Accordingly, inhibition of JNK signaling by expression of *Bsk*^DN^ significantly reduced the upregulation of *dilp8* mRNA levels observed upon RA activation (Figure 5B). Similarly, Egr/JNK pathway inhibition suppressed RA-induced Dilp8-GFP expression (Figure 5E). To examine whether Dilp8 production following RA^CS^ expression is required to reduce growth of adjacent cell populations, we expressed *dilp8*^RNAi^ along with RA in wing discs using *en-GAL4* and analyzed adult wing size. As shown in Figure 5F, depletion of *dilp8* totally rescued the non-autonomous reduction in tissue size caused by RA^CS^ expression. Interestingly, RA expression in the eye discs (*ey*>RA^CS^) also showed a non-autonomous reduction of adult wing size, indicative of inter-organ growth coordination (Figure 5G). Notably, RNAi-mediated inhibition of Grnd or Dilp8 in slow-growing eye discs fully rescued wing size (Figure 5G). We next analyzed a possible role of Dilp8 in RA-induced non-autonomous reduction of proliferation rates. Surprisingly, Dilp8 inhibition was not able to rescue the decreased BrdU incorporation levels caused by RA^CS^ or *dmyc*^RNAi^ expression in the unperturbed domain (Figures 5H-K). Altogether, these results indicate that dILP8 expression downstream of Egr/JNK is required to coordinate organ growth and final tissue size, but is dispensable for cell cycle adjustment.

**Figure 5:**
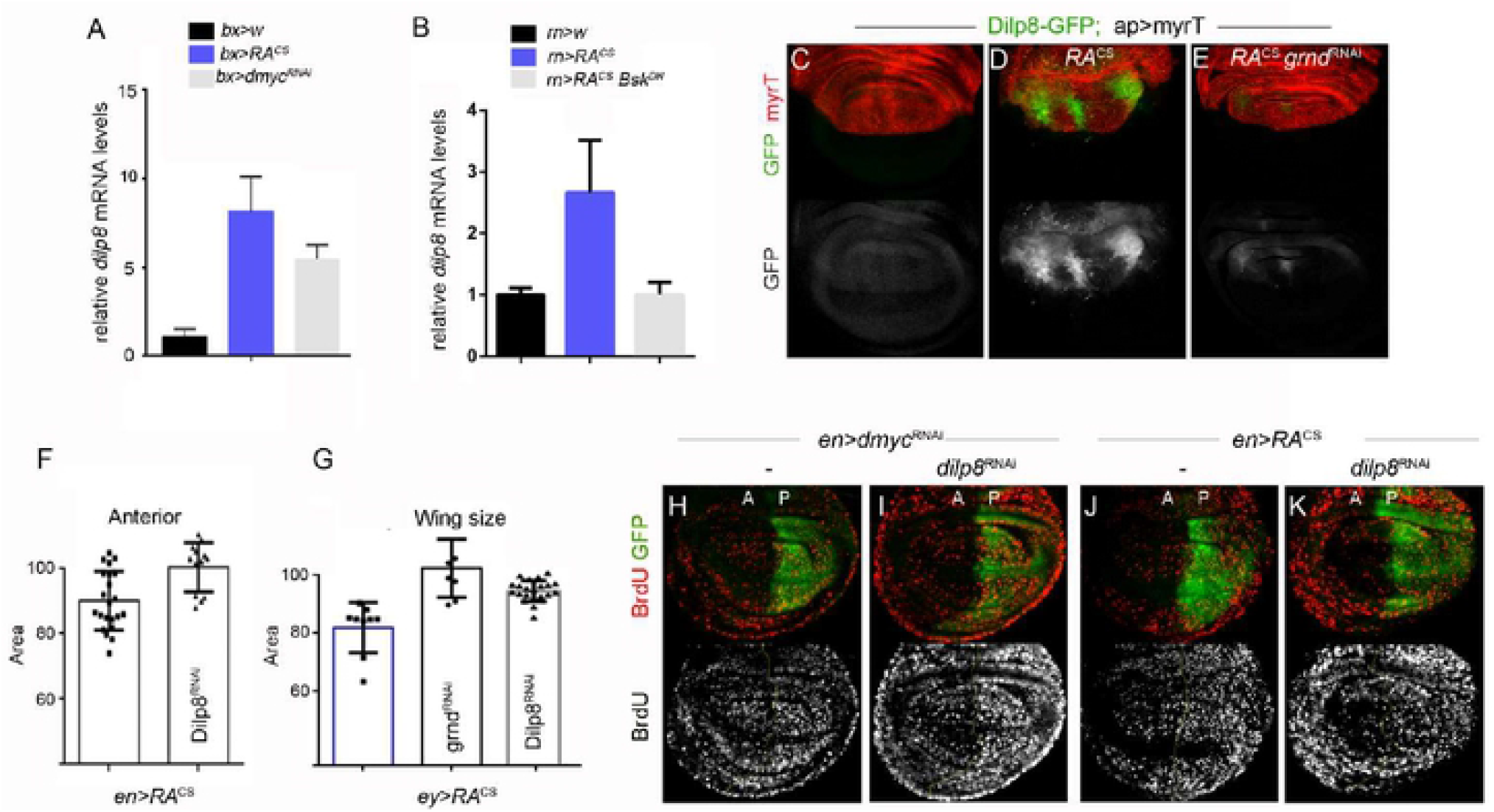
JNK-dependent Dilp8 expression contributes to non-autonomously reducing tissue size. (A-B) qRT-PCR showing *dilp8* transcript levels in wing discs (A) or full larvae (B) expressing the indicated transgenes under the control of *bx*- or *rn-Gal4*. Results are expressed as fold induction respect to control wing discs. (C-E) Wing discs carrying a Dilp8-GFP protein trap and expressing the indicated transgenes with *ap-Gal4* (expressed in dorsal compartment) were labeled to visualize GFP (green or grey) and myrTomato (red). (F) Histogram plotting normalized area of the A compartment of adult wings from individuals expressing RA^CS^ alone or in combination with *dilp8*^RNAi^ using *en-Gal4*. (G) Histogram plotting normalized wing size from individuals expressing RA^CS^ along with the indicated transgenes with *ey-Gal4* (drives expression to the eye imaginal disc). (H-K) BrdU incorporation assay in larval wing discs from individuals expressing GFP along with the indicated transgenes under the control of *en-Gal4*. Note the non-autonomous reduction in BrdU incorporation levels caused by *dmyc*^RNAi^ and RA^CS^ expression in Dilp8 knockdown wing discs.

### Caspase-dependent ROS production is required to non-cell-autonomously reduce proliferation rates downstream of Eiger/JNK signaling

Our GO analysis revealed an enrichment in RA^CS^-expressing wing discs of genes involved in cell redox homeostasis including expression of several glutathione S-transferases and other detoxifying genes (Figure 1H and Table S3), suggesting changes in the levels of Reactive Oxygen Species (ROS). To monitor ROS production in developing wing primordia we used the *gstD1*-GFP reporter, which is activated by ROS in *Drosophila* tissues [46], and observed that GFP expression was strongly induced in both the autonomous and non-autonomous territories following RA^CS^ or *dmyc*^RNAi^ expression (Figure 6A-D). Notably, whereas *gstD1*-GFP expression was mainly observed at the basal side of the epithelium in transgene-expressing compartment (Figure 6D’), consistent with cellular stress and apoptosis induction in the autonomous territory, *gstD1*-GFP expression in the non-autonomous territory was observed in intact epithelial cells (Figure 6D). These observations might suggest that ROS produced in the slow growing cell population trigger an antioxidant response in the neighboring compartment.

**Figure 6:**
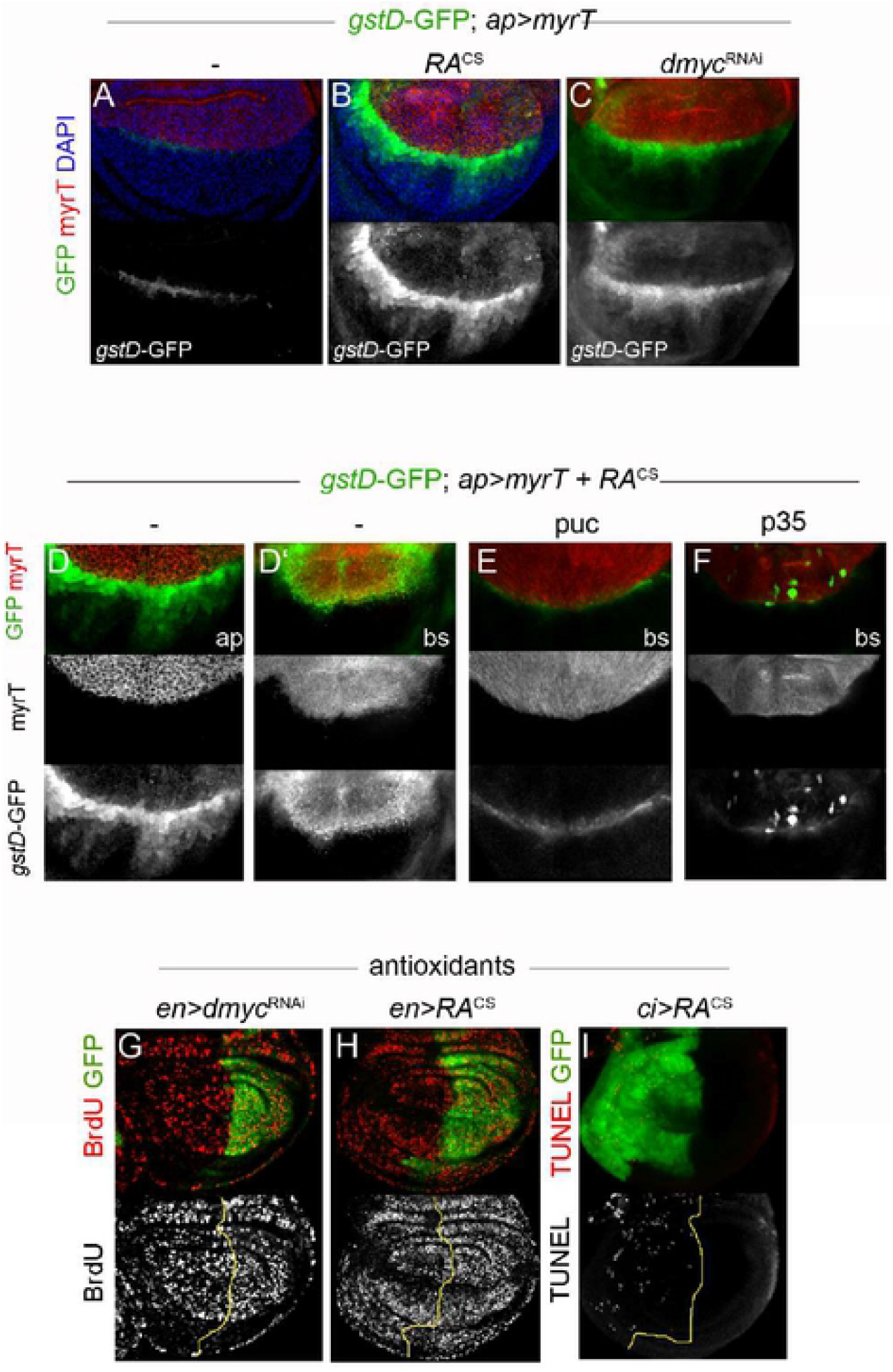
JNK and Caspase-dependent ROS production contribute to non-autonomously reduce proliferation rates. (A-F) Wing discs carrying the *gstD1*-GFP reporter and expressing the indicated transgenes with *ap-Gal4* were labeled to visualize GFP (green or grey), myrTomato (red) and DAPI (blue). RA^CS^ and *dmyc*^RNAi^ expression showed increased levels of *gstD1*-GFP in both the transgene-expressing and non-expressing domains (B-D). Co-expression of Puckered (puc; E) or p35 (F) fully rescued RA-induced *gstD1*-GFP levels. ap, apical; bs, basal. (G-I) BrdU and TUNEL assays in larval wing discs from individuals expressing GFP along with RA^CS^ or *dmyc*^RNAi^ under the control of *en*- or *ci-Gal4* and cultured in a medium supplemented with anti-oxidants (N-acetylcysteine, vitamin C and vitamin E).

ROS are associated with cellular stress and tissue damage, and provide important signaling during wound healing and regeneration in many model organisms [47–49]. Interestingly, during *Drosophila* wing regeneration dying cells generate a burst of ROS that propagate to the nearby surviving cells stimulating JNK signaling which is required for tissue repair [50]. We then asked whether ROS were produced downstream of the apoptotic pathway in slow growing compartments. To address this, we inhibited apoptosis in RA^CS^-expressing cells and assessed levels of *gstD1*-GFP expression. Notably, inhibiting apoptosis by expressing the baculovirus caspase inhibitor p35 largely impaired RA-induced *gstD1*-GFP expression in both transgene-expressing and adjacent wild type territories (Figure 6F). Consistently with a role of Egr/JNK signaling upstream of the apoptotic pathway, *gstD1*-GFP was not induced upon expression of the JNK phosphatase Puckered or in homozygous *egr^3^* wing discs (Figure 6E and data not shown).

We have previously shown that activation of caspase apoptotic pathway in slow growing domain is required to reduce proliferation rates of adjacent cell populations [17]. Interestingly, the non-autonomous reduction of BrdU incorporation levels caused by RA^CS^ or *dmyc*^RNAi^ expression was largely rescued by supplementing the medium with anti-oxidants N-acetylcysteine (NAC), vitamin C and vitamin E (Figure 6G-H). Importantly, reducing ROS levels by supplementing the medium with antioxidants did not rescue RA-induced apoptosis (Figures 6I). Similarly, Dilp8-GFP expression levels were still upregulated upon RA^CS^ or *dmyc*^RNAi^ expression when reducing ROS levels (data not shown). Collectively, these results indicate that ROS production downstream of Egr/JNK and the apoptotic machinery contributes to the non-autonomous reduction in proliferation rates.

## DISCUSSION

Intra- and inter-organ growth coordination is essential during animal development to buffer local growth perturbations and to produce individuals of proper size and shape. Despite its relevance, molecular mechanisms underlying this process are poorly understood. Using the *Drosophila* wing as a model system we have previously demonstrated that impairing growth within defined territories along the wing primordium triggers a Dmp53-dependent reduction of growth in adjacent non-perturbed cell populations [17]. Here, we have provided evidence that Dmp53 regulates intra-organ growth by inducing expression of the *Drosophila* TNF ligand Eiger [22,23]. We showed that Eiger-dependent activation of JNK pathway in slow growing compartments is required to non-cell-autonomously reduce growth and proliferation of adjacent cell populations. Furthermore, our work implies that tissue growth and proliferation rates can be uncoupled and are independently regulated by two different molecular mechanisms downstream of Eiger-JNK signaling (Figure 7). On one hand, Eiger/JNK-dependent dILP8 expression is required to systemically coordinate organ growth and final tissue size and dispensable for cell cycle adjustment. On the other hand, ROS production downstream of Eiger/JNK acts locally to regulate proliferation rates of adjacent epithelial cells. In this way, we show how signals coming either from neighboring cells or produced at a systemic level can orchestrate to accomplish growth coordination among different parts of an organ in a non-autonomous manner (Figure 7).

**Figure 7:**
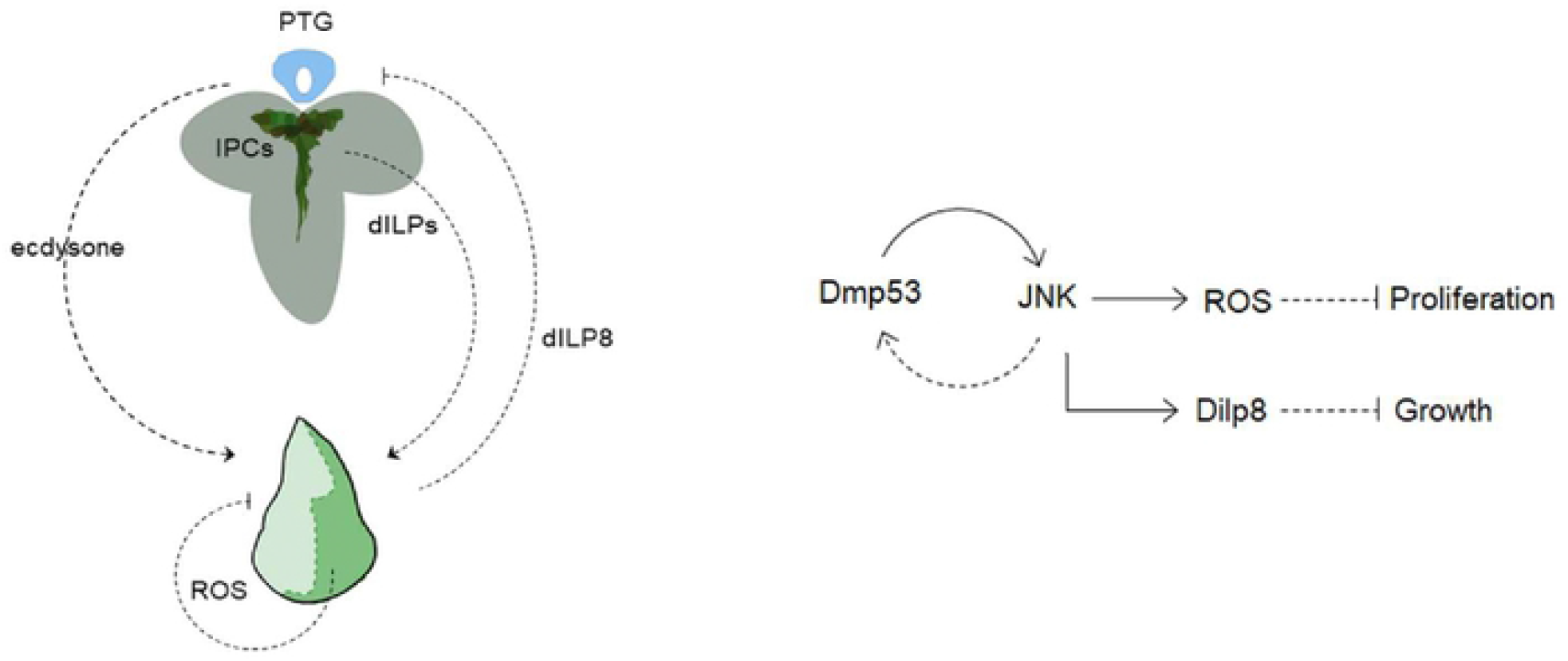
A Dmp53-mediated two step mechanism regulates growth and proliferation in adjacent populations through the production of systemic and local signals. Targeted expression of RA or depletion of dMyc activates Eiger-JNK signaling downstream of Dmp53. JNK-dependent expression of Dilp8 plays a crucial role in systemically coordinating the size of adjacent cell populations. On the other hand, JNK-dependent ROS production acts in a tissue local manner to regulates proliferation rates in adjacent cell populations.

TNF family ligands are well conserved throughout evolution acting mainly through JNK signaling to regulate growth, proliferation and apoptosis [51,52]. *Drosophila* type II transmembrane protein Eiger belongs to this family and plays an intrinsic tumor suppressor activity in epithelia by eliminating oncogenic cells through local endocytic JNK activation [53]. Interestingly, Eiger expression in apoptotic cells activates JNK pathway in neighboring cells thus propagating apoptotic JNK signaling along the tissue as part of a process called apoptosis-induced apoptosis (AiA) [28]. Moreover, it has been reported that interactions between tumor cells and tumor microenvironment mediated by Eiger and its receptor Grnd are as well able to drive JNK activation, tumor growth, and invasive behavior [53–55]. In this work, we showed that Eiger expression in growth deficient cell populations is required for both apoptotic and proliferative responses in unperturbed territories. However, Eiger-dependent JNK activation was restricted to slow growing domains and inhibition of Grnd-JNK signaling in the autonomous territory was sufficient to restore normal size in adjacent domains. Therefore, even though Eiger can act as a soluble ligand and activate JNK signaling in distant cells under certain circumstances, the role of Eiger in coordinating intra-organ growth seems to be restricted to the autonomous tissue and involves local JNK activation.

As mentioned before, JNK pathway is well known to control proliferation and apoptosis during regeneration and tumorigenesis in *Drosophila* imaginal discs [33–36,56]. In addition, JNK signaling regulates organ size during development through a non-canonical, Jun/Fos-independent mechanism [37]. Here we provided evidences that JNK coordinates intra-organ growth in a Jun/Fos-dependent manner through a mechanism involving expression of Dilp8 and generation of ROS, two different signaling events that independently regulate tissue size and cell number. Dilp8 has been recently identified as a signaling molecule produced by damaged imaginal discs able to delay metamorphosis and systemically reduce growth [39,40,57]. In developing wing discs, we have showed that Dilp8 expression in slow growing cell populations contribute to size reduction of adjacent wild-type territories, thereby contributing to organ proportion maintenance. Production of dILP8 by stressed or growth defective tissues activates Lgr3+neurons in the central brain to control synthesis of molting hormone ecdysone and/or insulin-like peptides, therefore coordinating growth with developmental timing [43,54,58]. In addition to its role in controlling developmental transitions, moderate ecdysone signaling has been shown to promote growth of imaginal discs both *in vitro* and *in vivo* [59–62]. Indeed, organ growth coordination in response to localized growth defects has been shown to be mediated, at least in part, by reduced ecdysone levels [19,21]. Remarkable, coordination of growth among adjacent wing disc compartments can be disrupted by exogenously feeding 20-hydroxyecdysone (20E) [19]. Therefore, Eiger/JNK-induced Dilp8 expression in slow growing compartment most likely regulates growth of adjacent unperturbed territory via modulation of ecdysone levels.

In addition to its function in driving apoptosis, reactive oxygen species (ROS) also act as signaling molecules influencing numerous cellular responses including cell proliferation, differentiation and senescence [63–65]. Although p53 is an important regulator of intracellular ROS levels in vertebrates, downstream mediators of p53 remain to be elucidated [66]. Here we have shown that ROS production, downstream of JNK and the apoptotic machinery, is required to reduce non-cell-autonomously proliferation rates in *Drosophila* developing epithelial tissues. In recent reports, ROS were found to be required for apoptosis-induced proliferation (AiP) and tissue regeneration of *Drosophila* imaginal discs [50,67]. In the undead model of AiP, extracellular ROS generated by the NADPH oxidase Duox recruits hemocytes to undead tissue. Hemocytes release Eiger, which promotes JNK activation in epithelial disc cells driving compensatory proliferation and hyperplastic growth of the tissue [67]. We found no evidence that hemocytes have this role in coordinated intra-organ growth, strongly suggesting that ROS-dependent hemocytes recruitment is context dependent and likely rely on sustained signals produced by undead cells. Following genetic or physical tissue damage, apoptotic cells generate a burst of ROS that propagate to the nearby surviving cells stimulating JNK signaling which is required for cell proliferation and tissue repair [50]. Thus, whereas ROS-stimulated JNK activity promotes cell proliferation in the context of tissue regeneration, JNK-induced ROS non-cell-autonomously reduces proliferation to adjust cell number.

## MATERIALS AND METHODS

### *Drosophila* strains and maintenance

The following *Drosophila* stocks were used: *UAS-Ricin^CS^* (BL38623), *en-Gal4* (BL1973), *bx-Gal4*(BL8860), *UAS-Basket-DN* (BL6409), *UAS-p35* (BL6298), *UAS-Dmp53^H159N^* (BL8420), *UAS-Dmp53* (BL8418), *UAS-Fos^C-Ala^* (UAS-Fos^DN^ in the text [33]), *UAS-Tak1^K46R^* (Tak^DN^ in the text (BL58811)), hep^r75^ (BL6761), *UAS-eiger*^IR^ [22], *UAS-eiger^RNAi^* (VDRC108814), *UAS-grindewald*^RNAi^ (VDRC104538); *UAS-dmyc*^RNAi^, *UAS-Dmp53*^RNAi^ (VDRC38235), *eiger^1^* and *eiger^3^* [22], *eiger-lacZ* [30], *Eiger-GFP* (fTRG library; VDRC318615). Other stocks are described in Flybase. RNAi lines for secreted signaling molecules were obtained from Vienna *Drosophila* RNAi Center (VDRC) (Supporting Information Table S2).

Flies were reared at 25°C on *Drosophila* standard medium (4% glucose, 40 g/L yeast, 1% agar, 25 g/L wheat flour, 25g/L cornflour, 4 ml/L propionic acid and 1,1 g/L nipagin).

For experiments using the cold sensitive version of Ricin-A (RA^CS^), embryos containing the Gal4 driver and the UAS-RA^CS^ transgene were collected for 24 h and maintain at 18°C (restrictive temperature). Early second instar larvae were switched to 29°C (permissive temperature) until late L3 (for wing disc experiments) or until adulthood (for adult wing analysis).

### Immunostaining

Third instar larvae were dissected in cold phosphate-buffered saline (PBS) and fixed in 4% formaldehyde/PBS for 20 min at room temperature. Dissected larvae were then washed and permeabilized in PBT (0,2% Triton X-100 in PBS) for 30 min and blocked in BBT (0,3% BSA, 250mM NaCl in PBT) for 1 hour. Samples were incubated with primary antibody diluted in BBT overnight at 4°C, washed three times (15 min each) in BBT and incubated with secondary antibodies and DAPI (1μg/ml) for 1,5 hour at room temperature. After three washes with PBT (15 min each), wing discs were mounted in mounting medium (80% glycerol/PBS containing 0.05% n-Propyl-Gallate). All steps were performed on a rocking platform at the indicated temperature. The following primary antibodies were used: mouse anti-BrdU (G3G4; Developmental Studies Hybridoma Bank (DSHB)), mouse anti-MMP1 (3A6B4, DSHB), mouse anti-p53 (7A4, DSHB), rabbit anti-p-Histone H3 (sc-8656, Santa Cruz), rabbit anti-β-Gal (A11132, Invitrogen), sheep anti-Digoxigenin-AP (#11093274910, Roche). The secondary antibodies used were: anti-mouse IgG-Alexa Fluor 594, anti-mouse IgG-Alexa Fluor 488, anti-rabbit IgG-Alexa Fluor 594 and anti-rabbit IgG-Alexa Fluor 488 (Jackson InmunoResearch).

### TUNEL and BrdU assays

TUNEL was performed as described in [68] using In Situ Cell Death Detection Kit provided by Roche Diagnostics. BrdU incorporation was performed as described in [68]. Briefly, third-instar larvae were dissected in PBS and incubated with 5-bromo-2’-deoxy-uridine (10 μM, Roche) for 45 min, washed and fixed in 4% formaldehyde. Following incubation with HCl (2N) for 30 min samples were neutralized with Borax (100 mM) and immunostained as before.

### Image processing

Images were acquired on a Leica SP8 inverted confocal microscope. Images were analyzed and processed using Fiji [69] and Adobe Photoshop. In the images presented, wing discs orientation and/or position were adjusted in the field of view. No relevant information has been affected. The original images are available on request.

### RNA in situ hybridization

In situ hybridization was performed as described in [68]. Antisense DIG-labelled *eiger* RNA probe was prepared from XhoI-linearized pBSK-eiger plasmid using T3 RNA polymerase (DIG RNA Labeling Kit, Roche) and detected with the DIG Nucleic Acid Detection Kit (Roche). Wing imaginal discs were mounted in glycerol and imaged with a Nikon E200 bright-field microscope.

### Quantification of adult wing size and cell number

Size of the A and P compartments in adult wings was measured using Fiji. Cell density was measured as the number of hairs (each wing cell differentiates a hair) per defined area as previously described [17]. At least 10 wings per genotype were scored. Calculated area and cell density values for the different genotypes were normalized to control GFP-expressing wings. Average values and corresponding standard deviation (SD) were calculated and two-tailed unpaired student’s t test was carried out. Calculations and bar graphs were made using the Graph pad Prism 7 software.

### RNA isolation and quantitative RT-PCR

For quantification of mRNA levels, total RNA was extracted from wing imaginal discs of 30 larvae using TRIZOL RNA Isolation Reagent (Invitrogen). First strand cDNA synthesis was performed using an oligo(dT)18 primer and RevertAid reverse transcriptase (Thermo Scientific) under standard conditions. Quantitative PCR was performed on an aliquot of the cDNA with specific primers (Supplementary Table 4) using the StepOnePlus Real-Time PCR System. Expression values were normalized to actin transcript levels. In all cases, three independent samples were collected from each condition and genotype, and duplicate measurements were conducted.

### Chromatin immunoprecipitation

ChIP assays were performed according to modEncode protocol [70]. Fifty L3 larvae were dissected for ChIP assays using anti-p53 antibodies (DSHB; 7A4). Primer used to detect immunoprecipitated DNA are listed in Supplementary Table 4.

### Microarray

Transcription profiles were performed at the Functional Genomics Core of IRB as previously described [71]. Three independent RNA samples were prepared from wing imaginal discs of late L3 larvae expressing GFP or GFP plus RA^CS^ under the control of en-gal4. After dissection of wing discs in cold PBS, the posterior GFP-positive compartment was separated under a fluorescence stereoscope and used for RNA extraction with Trizol. Library preparation and amplification were performed using Ovation RNA Amp System V” (Nugen;3100-60). Fragmentation and labelling were performed using Encode Biotin Module (Nugen; 4200-60). Hybridization mixture was prepared following the Affymetrix protocol. Each sample was hybridized to a GeneChip Drosophila Genome 2.0 Array (Affymetrix), washed and stained in a GeneChip Fluidics Station 450 (Affymetrix). Microarray scanning and CEL files generation were performed using an Affymetrix GeneChip Scanner GSC3000. To generate log2 expression estimates, overall array intensity was normalized between arrays and the probe intensity of all probes in a probe set summarized to a single value using gcRMA (gcRMA 2.0.0; Bioconductor package). The transcriptional profile of RA expressing and nonexpressing cells of the posterior compartment was compared and those genes with fold changes ≥ or ≤ 1 and p value ≤0,05 were considered to be differentially expressed.

Gene ontology (GO) enrichment analysis of those transcripts that were significantly upregulated or downregulated was performed using DAVID v6.7. Microarray datasets have been deposited in the Gene Expression Omnibus (http://www.ncbi.nlm.nih.gov/geo/) under accession number GSE125794.

## Acknowledgements

We thank Daniel Gonzalez, Pablo Wappner, Fernanda Ceriani and Andrés Garelli for support and reagents, the Developmental Studies Hybridoma Bank, Vienna Drosophila Resource Center (VDRC) and Bloomington Stock Center for antibodies and fly strains. We thank Agustin Arce for help in microarray analysis, and Leandro Lucero for guidance in ChIP experiments. J.A.S and M.C.I are funded PhD fellowships from CONICET. F.A. is a member of CONICET. M.M. is an ICREA Research Professor, and M.M.’s lab is funded by the SIGNAGROWTH-BFU2013-44485 and INTERGROWTH-BFU2016-77587-P grants from the Ministerio de Ciencia, Innovación y Universidades (Government of Spain), and ERDF “Una manera de hacer Europa”. A.D. is a member of CONICET and Professor at UNL. This work was supported by grants from the Agencia Nacional de Promoción Científica y Tecnológica, Argentina (ANPCyT), Universidad Nacional del Litoral (UNL) and MinCyT-DAAD Bilateral Cooperation Program.

## Author Contributions

J.A.S. performed experiments and analyzed data. D.M. performed experiments. M.C.I. performed qRT-PCRs and edited the manuscript. F.A. contributed to ChIP experiments. M.M. participated in experimental design, analyzed data and edited the manuscript. A.D. designed experiments, analyzed data, and wrote the manuscript.

## Supporting Information Legends

**Supplementary Figure S1**

(A) qRT-PCR showing transcript levels of a selected group of genes in wing discs expressing *RA^CS^* or *myc^RNAi^* with *en-Gal4* or *Bx-Gal4*. Results are expressed as fold induction relative to control wing discs. (B) Signaling pathways affected in RA expressing wing discs. (C) Venn diagrams showing overlap between differentially expressed genes in RA-expressing cells and previously identified p53 target genes (Akdemir et al., 2007; van Bergeijk et al., 2012).

**Supplementary Figure S2**

(A) List of genes coding for extracellular proteins that were differentially expressed in RA-expressing cells and corresponding fold change. (B) Histogram plotting normalized area of the anterior (A) compartment of adult wings from individuals expressing RA with hh-Gal4 in eiger^3/3^ mutants. ** p<0.01. (C) Wing imaginal discs from individuals expressing RA^CS^ along with eiger^RNAi^ under the control of *en-gal4* and stained to visualize PH3 levels.

**Supplementary Figure S3**

(A) Expression pattern of egr-lacZ (Muzzopappa et al., 2017) and Eiger-GFP (Muzzopappa et al., 2017; Sarov et al., 2016) reporters in the eye and wing imaginal discs of wild type larvae. (B) Wing discs carrying Eiger-GFP protein trap and stained to visualize GFP (green). Myc^RNAi^-expressing cells displayed increased levels of Eiger-GFP. (C) qRT-PCR plotting *corp* mRNA levels in wing discs expressing *RA^CS^* or *RA^CS^* plus *Dmp53^H159.N^* with en-Gal4. Results are expressed as fold induction respect to control wing discs (en>w).

**Supplementary Figure S4**

(A-B) qRT-PCR showing *mmp1* and *puc* mRNA levels in wing discs expressing the indicated transgenes under the control of en-Gal4 relative to control wing discs (en>w).

**Supplementary Table 1:**List of genes differentially expressed between RA-expressing and wild type cells

**Supplementary Table 2:**List of genes encoding for known or predicted secreted molecules and their corresponding transgenic RNAi lines used for the screen

**Supplementary Table 3:**Signaling pathways affected in RA-expressing cells

**Supplementary Table 4:**List of primers used in this study

